# A new mechanism for tree mortality due to drought and heatwaves

**DOI:** 10.1101/531632

**Authors:** Hervé Cochard

**Affiliations:** Université Clermont-Auvergne, INRAE, PIAF, 63000 Clermont-Ferrand – France

**Keywords:** Water stress, climate change, forest, dieback, ecophysiology, temperature

## Abstract

Plants tend to die earlier in hot and drought conditions, but the underlying mechanisms are not yet understood. I propose here a new mechanism by which excessive residual water losses caused by high cuticular permeabilities and a high leaf-to-air vapor pressure deficits would trigger uncontrolled and sudden cavitation events. The combination of heat and drought stresses may therefore lead to an unsuspected risk of hydraulic failure. I explored this hypothesis with a new mechanistic model. The simulations support this hypothesis and highlight the critical role played by the cuticle phase transition temperature. Experiments are now awaited to confirm these predictions.

## Introduction

In recent years, cases of widespread forest mortality have been recorded worldwide (Allen *et al* 2010). These die-offs seem mostly triggered by extreme drought and heat stresses (Anderreg *et al* 2016, Adams *et al* 2017a, Williams *et al* 2013). Current models predict an increased risk of mortality due to climate warming (McDowell and Allen 2015). However, the exact physiological mechanisms responsible for tree mortality under these extreme events remain largely unknown, which hinders our ability to model and predict the risk of forest dieback and changes in species distribution ranges.

Drought-induced tree mortality is tightly associated with the risk of hydraulic failure, whereby excessive tensions in the xylem tissue provoke cavitation events that block water transport from the roots to the leaves. Although most species live on the verge of hydraulic failure (Choat *et al* 2012), trees seem remarkably protected against xylem dysfunctions during drought events (Cochard and Delzon 2013) suggesting that drought-induced hydraulic failure may occur only under extreme and peculiar climatic conditions.

There is a large body of observations and experimental evidences suggesting that heatwaves exacerbate the risk of mortality under drought conditions (Adams *et al* 2017a). This was particularly noticeable during the 2003 hot-drought in France (Landmann and Dreyer 2006). A temperature-driven carbon starvation caused by an increase in respiration costs has been proposed as a mechanism for tree die-off under hot droughts (Adams *et al* 2009). However, more recent studies suggest that trees are rarely under the threat of carbon starvation (Adams *et al* 2017b).

Here, I explore a new hypothesis for tree mortality under extreme heat and drought stresses whereby tree hydraulic safety margin would be strongly reduced under these extreme conditions. To our knowledge, this hypothesis has not been formulated and evaluated to date. I tested this hypothesis with the mechanistic model SurEau (Martin St-Paul *et al* 2017, Cochard *et al* 2020) developed to simulate plant water relations and hydraulic functioning during water stress. My objective is not provide a full description of the model, which is available elsewhere (Cochard *et al* 2020), but to focus only on the key physiological processes temperature-dependent.

## Theory: Putative temperature effects on plant hydraulics

From a thermodynamic point of view, air temperature (*T*_*air*_) can impact tree hydraulics and water relations in several ways (figure 1).

**Figure 1:**
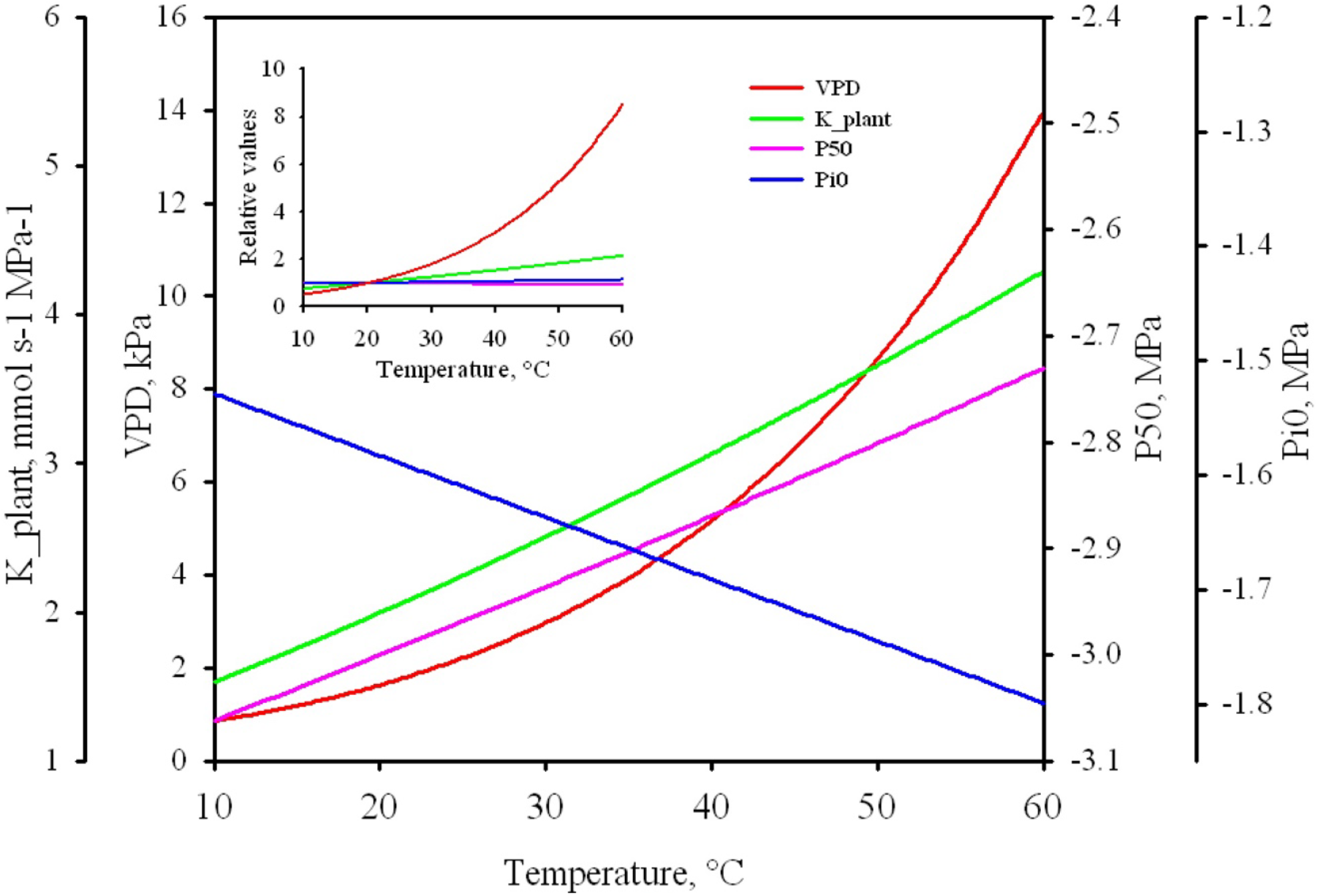
Theoretical effects of temperature on air *VPD* (red), plant hydraulic conductance *K*_*plant*_ (green), xylem vulnerability to cavitation *P*_*50*_ (pink) and osmotic potential Pi0 (blue). The relative variations of these parameters are shown in the insert.

The **vapor pressure deficit** (*VPD*) between the atmosphere and the leaf is the driving force for leaf water loss. A rise in air temperature strongly increases leaf transpiration via its exponential effect on air saturation vapor pressure. This microclimatic effect is exacerbated under drought conditions because stomatal closure decreases the rate of heat loss by evaporation which increases leaf temperature.

The *VPD* between the air and leaf surface is given by:

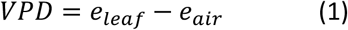

where *e*_*leaf*_ and *e*_*air*_ are, respectively, the vapor pressure (Pa) inside the leaf and bulk air levels. These values are dependent of the saturation vapor pressure that itself is a function of temperature according to Buck equation (Buck 1981):

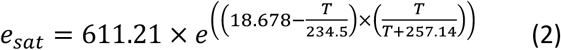

where *T* represents the leaf (*e*_*sat_leaf*_) and the air (*e*_*sat_air*_) temperature (°C). The actual vapor pressure at the leaf level is further decreased by water deficit of the leaf due to its negative water potential *Ψ*_*leaf*_ according to Nobel (2009):

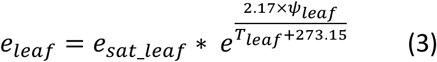

Similarly, the air vapor pressure is influenced by the air relative humidity *RH* (%):

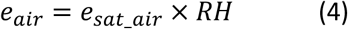

The effect of *T*_*air*_ on *VPD* is shown on figure 1 (red line) assuming RH=30% and *T*_*leaf*_ = *T*_*air*_.

Clearly, *T*_*air*_ has a strong and exponential effect on *VPD*, the effect been exacerbated if *T*_*leaf*_ is above *T*_*air*_ which frequently occurs when stomata are closed. Therefore, *T*_*air*_ has a strong effect on leaf transpiration *E* (mmol s^−1^ m^−2^) as:

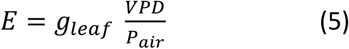

where *g*_*leaf*_ is the leaf conductance to vapor water (mmol s^−1^ m^−2^), and *P*_*air*_ the atmospheric pressure. *g*_*leaf*_ is composed of two resistances in parallel: the stomatal (*g*_*s*_) and the cuticular conductance (*g*_*min*_).

The **dynamic fluidity** *F*_*T*_ of liquid water, the reciprocal of its viscosity, varies with temperature according to the empirical formula derived from Lide (2004):

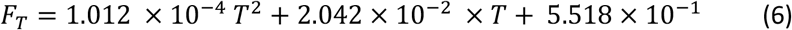

The hydraulic conductance (*K*_*T*_, mmol s^−1^ MPa^−1^) will vary linearly with *F* as:

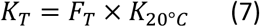

The effect of *T*_*air*_ on the total plant conductance *K*_*plant*_ will depend of the temperature of the different organs (root, trunk, shoots, leaves) and the conductance of each of these organs. In figure 1 (green line) and in the following simulations, I assumed that the root temperature was constant, the trunk and shoots temperatures were at equilibrium with *T*_*air*_, and the leaf temperature was function of its energy budget. I also assumed that 50% *K*_*plant*_ was above ground. The effect of *F*_*T*_ on *K*_*plant*_ is significant, with a roughly linear effect of 2.5% per °C for the range of temperature considered. Experimental data support this model (Cochard *et al* 2000).

The **surface tension** *σ*_*T*_ of liquid water against air decreases nearly linearly with temperature according to this empirical formula derived from Lide (2004):

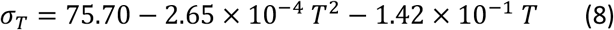

According to Young and Laplace equation, *σ*_*T*_ has a linear impact on the air-seeding xylem pressure triggering cavitation. I can then compute the effect of *T* on the mean *P*_*50*_ values of the different plant organs:

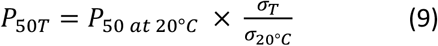

I can therefore predict a small increase (less negative) of *P*_*50*_ (about 0.2% per °C) in the range of temperature considered (figure 1, pink line), which is consistent with some experimental data (Cochard *et al* 2007)

The **osmotic potential** *Π*_*T*_ of leaf cells increases with leaf temperature according to van’t Hoff relation:

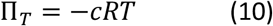

where *c* is the concentration of solutes in the cells and *R* the gas constant. Therefore, if *Π*_*20*_ is the osmotic potential at 20°C and *T* is expressed in °C I have:

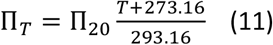

As the total leaf water potential *Ψ*_*leaf*_ remains constant when temperature is changing, the leaf turgor pressure *P*_*leaf*_ must also increase with temperature as:

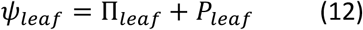

This may impact in return leaf stomatal conductance and transpiration under the hypothesis that these variables are determined by *P*_*leaf*_. Computations shows a linear and marginal decrease of *Π* with temperature at a rate of 0.34 % per °C (figure 1, blue line).

The **leaf cuticular conductance** to vapor water *g*_*min*_ is a parameter known to be strongly influenced by leaf temperature. Indeed, recent studies have demonstrated a sharp increase in *g*_*min*_ above a phase transition temperature *Tp* that matches the range of temperatures known to trigger mortality during hot-droughts (Riederer and Muller 2006; Schuster *et al* 2016). Following Riederer and Muller (2006) and Schuster *et al* (2016) I used a double Q_10_ equation to model the effect of *T*_*leaf*_ on *g*_*min*_ knowing *g*_*min_20*_ the values taken by *g*_*min*_ at 20°C and deriving *g*_*min_Tp*_ from the equation 12 below:
if *T*_*leaf*_ < *Tp* then

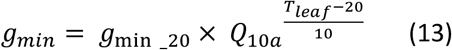

if *T*_*leaf*_ > *Tp* then

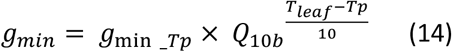

where *Q*_*10a*_ and *Q*_*10b*_ are the *Q*_*10*_ values of the relationship below and above *Tp*, respectively.

According to the experimental data of Riederer and Muller (2006), *Q*_*10a*_ and *Q*_*10b*_ appear relatively constant across species and equal to 1.2 and 4.8, respectively. By contrast, *Tp* seem a more variable parameter.

The figure 2 shows how *g*_*min*_ respond to *T*_*leaf*_ when *Q*_*10a*_ and *Q*_*10b*_ are constant but *Tp* allowed to vary between 30 and 45°C. Below *Tp*, *g*_*min*_ increases slightly with temperature. By contrast, above *Tp*, *g*_*min*_ increases sharply and exponentially with *T*_*leaf*_.

**Figure 2:**
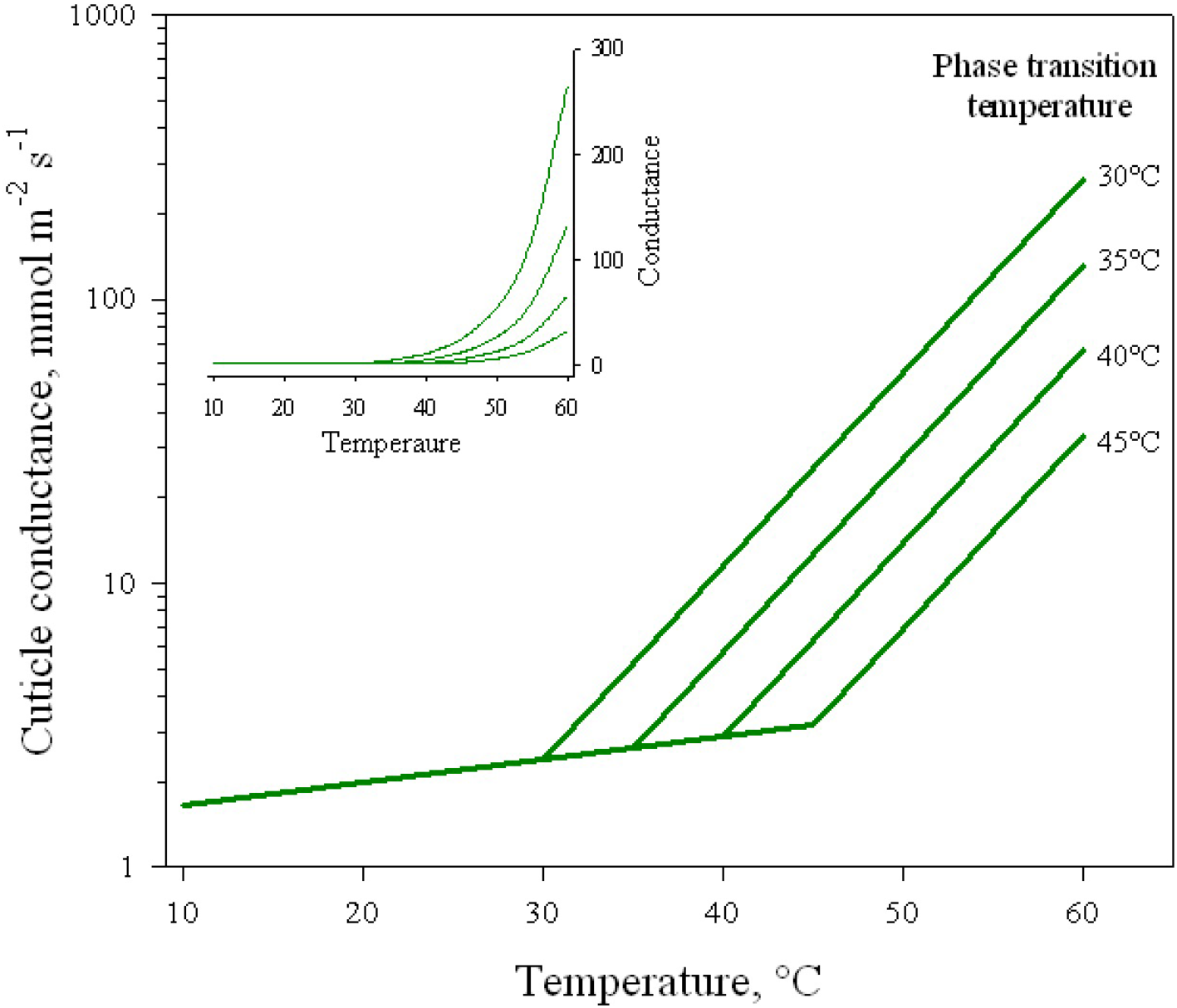
Bi-phasic model for the temperature dependence of leaf cuticular conductance (*g*_*min*_). The conductance follows two Q_10_ curves above and below a phase transition temperature *Tp.* The insert shows the curves on a linear *y* scale.

## Methods: Modeling the temperature effects on plant hydraulics

I used a new mechanistic model (SurEau, Martin-StPaul *et al* 2017, Cochard *et al* 2020) to predict the effect of air-temperature on the timing of hydraulic failure and putative mortality. To this aim, I developed a new version of SurEau_C, coded in the C-language, that dynamically model tree water relation, gas exchanges and hydraulics during water stress at a 1ms step. All the five processes described above were implemented in the model in addition to a leaf energy budget module that computes the leaf temperature of an illuminated and transpiring leaf. For that purpose I used the equations from classical text books (Jones 2013, Nobel 2009).

I simulated the drought response of a “typical” potted sapling, whose main characteristics are given in table 1. The results of the model are of course strongly determined by the initial conditions of the simulations. It is therefore necessary to pay more attention to the relative variations of the different simulated variables than to their absolute value.

**Table 1:** Main characteristics of the reference simulation use for the control plant.

| Parameters | Leaf area<br>m<sup>2</sup> | Plant Height<br>m | $g_{leaf}/g_{min}$<br>mmols<sup>-1</sup>m<sup>-2</sup> | $T_p$<br>°C | $Q_{10a}/Q_{10b}$ | $P_{50}$<br>MPa | Slope at $P_{50}$<br>%MPa<sup>-1</sup> | $\Pi_0$<br>MPa | $\varepsilon$<br>MPa | $K_{plant}$<br>mmols<sup>-1</sup>MPa<sup>-1</sup> |
| --- | --- | --- | --- | --- | --- | --- | --- | --- | --- | --- |
| Values | 0.5 | 1.3 | 165 / 2 | 35 | 1.2 / 4.8 | -3 | 100 | -1.58 | 7.55 | 4.4 |

The standard pedoclimatic conditions use for the simulations are given in table 2. Pedotransfer equations were used to compute soil water content and soil water potential (van Genuchten 1980).

**Table 2:** Main pedoclimatic conditions use for the control simulation.

| Parameters | Soil volume<br>m<sup>3</sup> | Soil type | $T_{air-min}$<br>°C | $T_{air-max}$<br>°C | $RH_{air-min}$<br>% | $RH_{air-max}$<br>% | PAR<br>μmol | Wind speed<br>ms<sup>-1</sup> |
| --- | --- | --- | --- | --- | --- | --- | --- | --- |
| Values | 0.05 | clay | 15 | 25 | 30 | 80 | 1500 | 1.0 |

The simulation started at midnight with a soil at full water holding capacity, let to dehydrate by evapotranspiration, and stopped at *t*_*HF*_ (days) when the level of embolism reached 99PLC, which was considered as the point of irreversible hydraulic failure. The values I report here for *t*_*HF*_, although realistic, should be taken as relative values because they are strongly dependent on the parameters of the simulations. For instance, doubling the *Soil volume* or dividing *Leaf area* by two will roughly double *t*_*HF*_.

I first tested the impact of the different mechanisms listed above on *t*_*HF*_ by varying *T*_*air-min*_ from 10 to 50°C and *T*_*air-max*_ from 20 to 60°C. I then simulated the impact of a heatwave by increasing *T*_*air*_ by +15°C for 3 consecutive days above the standard temperatures (15/25°C). The heatwave was set to start after 0 to 26 days after the onset of the simulation. I also conducted a sensitivity analysis of the parameters determining *g*_*min*_ by varying *T*_*p*_ from 30 to 40°C, *Q*_*10a*_ from 1.0 to 1.4 and *Q*_*10b*_ from 3.8 to 5.8, the heatwave starting at day 9. Finally, I analyzed the impact of increasing *Tp* from 25 to 55°C on minimum *t*_*HF*_ in response to heatwaves of +0 to +30°C occurring between days 0 to days 40.

## Results

### Temperature effects on hydraulic failure

In figure 3 I show the results of 3 simulations where midday air temperature *T*_*air*-max_ was set to 20, 25 and 30°C, respectively. I modelled only the effect of *T*_*air*_ on VPD in these simulations. Increasing *T*_*air*-max_ sharply increased the maximum midday transpiration rate (blue lines) because of its effect on *VPD*. As a consequence, plants exposed to higher temperatures depleted faster their soil water content and regulated their transpiration earlier. As a result, the time to stomatal closure (*t*_*g0*_, blue arrows) decreased with air temperature, with a 5 days difference between the two extreme simulations. As another consequence, cavitation was induced earlier for plants exposed to higher temperatures (red lines) and the time to hydraulic failure (*t*_*HF*_, red arrows) was strongly reduced by temperature (16 days difference between extreme temperatures). The effect of temperature on *t*_*HF*_ was more pronounced than on *t*_*g0*_, highlighting a critical role of VPD and the residual plant transpiration after stomatal closure on hydraulic failure. When *T*_*air*-max_ was allowed to increase to 60°C (figure 4, black line) *t*_*HF*_ further decreased following a negative exponential function.

**Figure 3:**
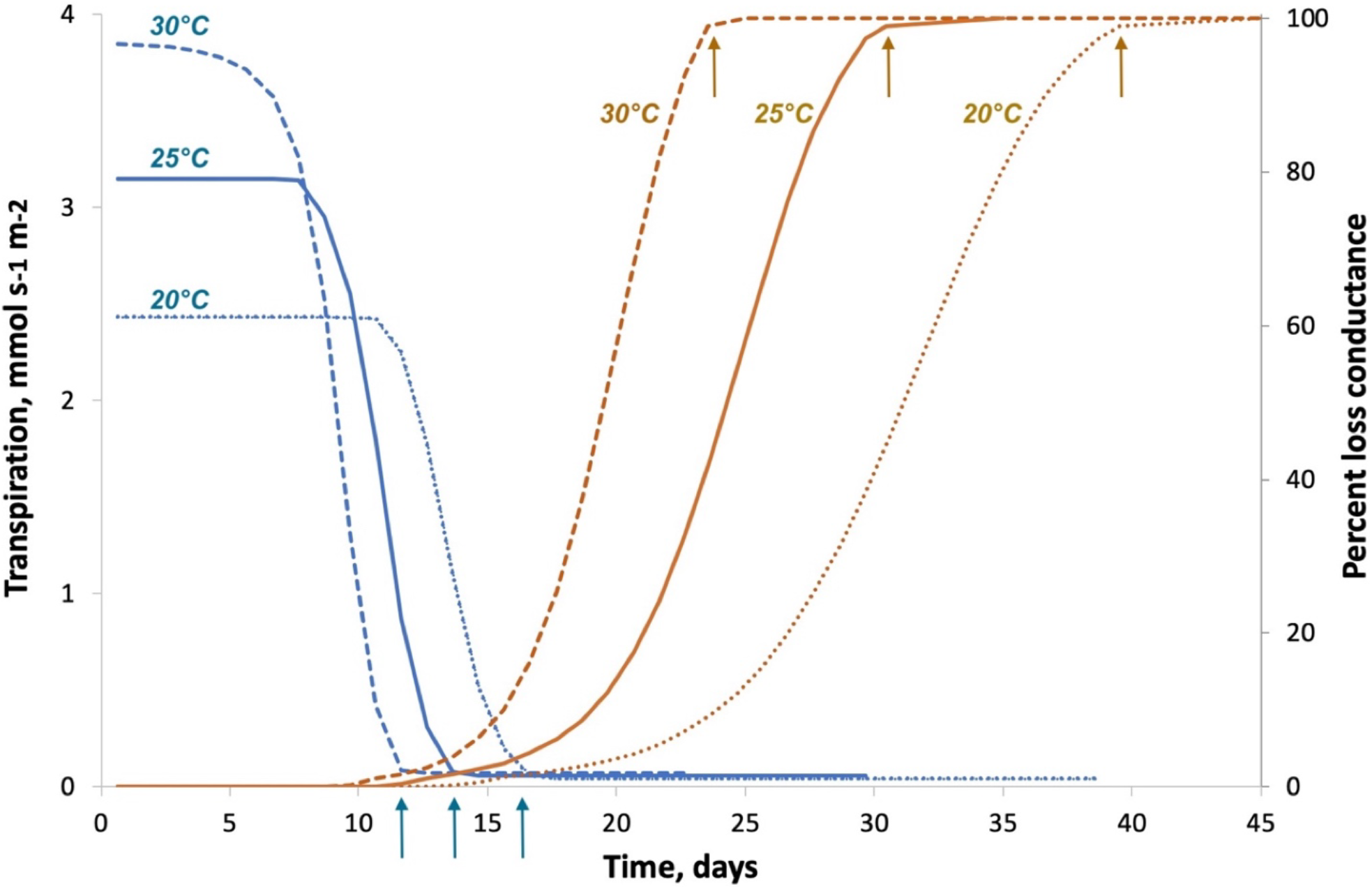
Typical time variations of leaf transpiration (blue) and xylem embolism (orange) for control plants exposed to a water stress at day 0 under three different air temperatures (different line types). Plants exposed to higher temperatures closed their stomata earlier (blue arrows) but displayed a shorter time to total hydraulic failure (orange arrows).

**Figure 4:**
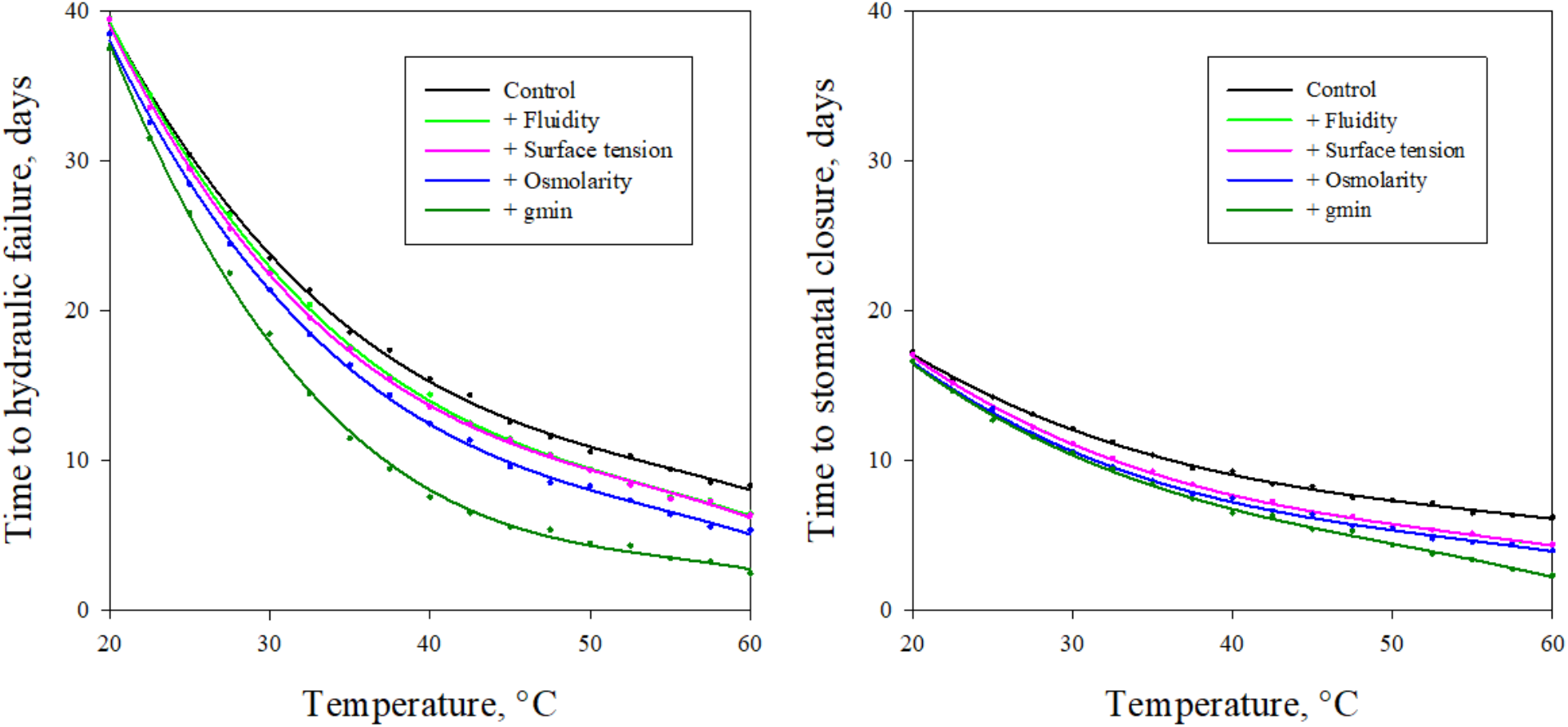
Impact of different processes thermodynamically determined by temperature on the timing of hydraulic failure (*t*_*HF*_) and stomatal closure (right). The black line represents the temperature response of *tHF* for a plant where only the VPD effect was modelled. The color lines show the additive effects of water fluidity (light green), water surface tension (pink), osmolarity (blue) and cuticular conductance (dark green).

In a second set of simulations, I added the impact of temperature on water fluidity which further decreased *t*_*HF*_ of about 1 day (figure 4, bright green line). This decreased was explained by a slight increase in transpiration rate caused by an enhanced hydraulic efficiency with temperature (higher *K*_*plant*_). The effect of temperature water surface tension was only marginal and the impact on *t*_*HF*_ hence negligible for the range of temperature considered (figure 4, pink line). Adding the effect of temperature on osmotic potential in the model decreased *t*_*HF*_ of roughly one more day (figure 4, blue line). This was explained by the effect of *Π*_*leaf*_ on leaf turgor *P*_*leaf*_ and leaf transpiration.

In a last set of simulations I further included a temperature dependence of *g*_*min*_ in the model according to the biphasic equations described above and setting the phase transition temperature *Tp* to 35°C. The model shows a sharp impact on *t*_*HF*_ that was further reduced by 5 days (figure 4, dark green line). The effect was noticeable even for temperatures below for temperature below *Tp* indicating a strong impact of even small variations of *g*_*min*_ on *t*_*HF*_.

The model suggests two major effects of air temperature on plant hydraulic failure. The first one, already well documented, is its impact on *VPD*, and hence transpiration and soil water use. However, the simulations highlight the critical role of residual leaf transpiration and its temperature dependence on hydraulic failure (Duursma *et al* 2018). The second major effect is related to the temperature dependence of *g*_*min*_, which also determine the residual transpiration rate. The time to hydraulic failure appears clearly more determined by the water losses beyond the point of stomatal closure rather than the speed at which plant empty the soil water reserve when stomata are still open.

### The impact of heatwaves on hydraulic failure

I simulated the impact of a +15°C heatwave lasting for 3 consecutive days and occurring at different times after the onset of a drought episode on the induction of cavitation (figure 5). The drought response of control a plant not exposed to a heatwave is shown as a black line, together with its transpiration rate (dotted pink line). Under the condition of this simulation, stomata started to close after 7 days and were completely closed after 13 days. At this stage, the degree of embolism was still less than 5% and progressively increased to reach >99PLC after 26 days.

**Figure 5:**
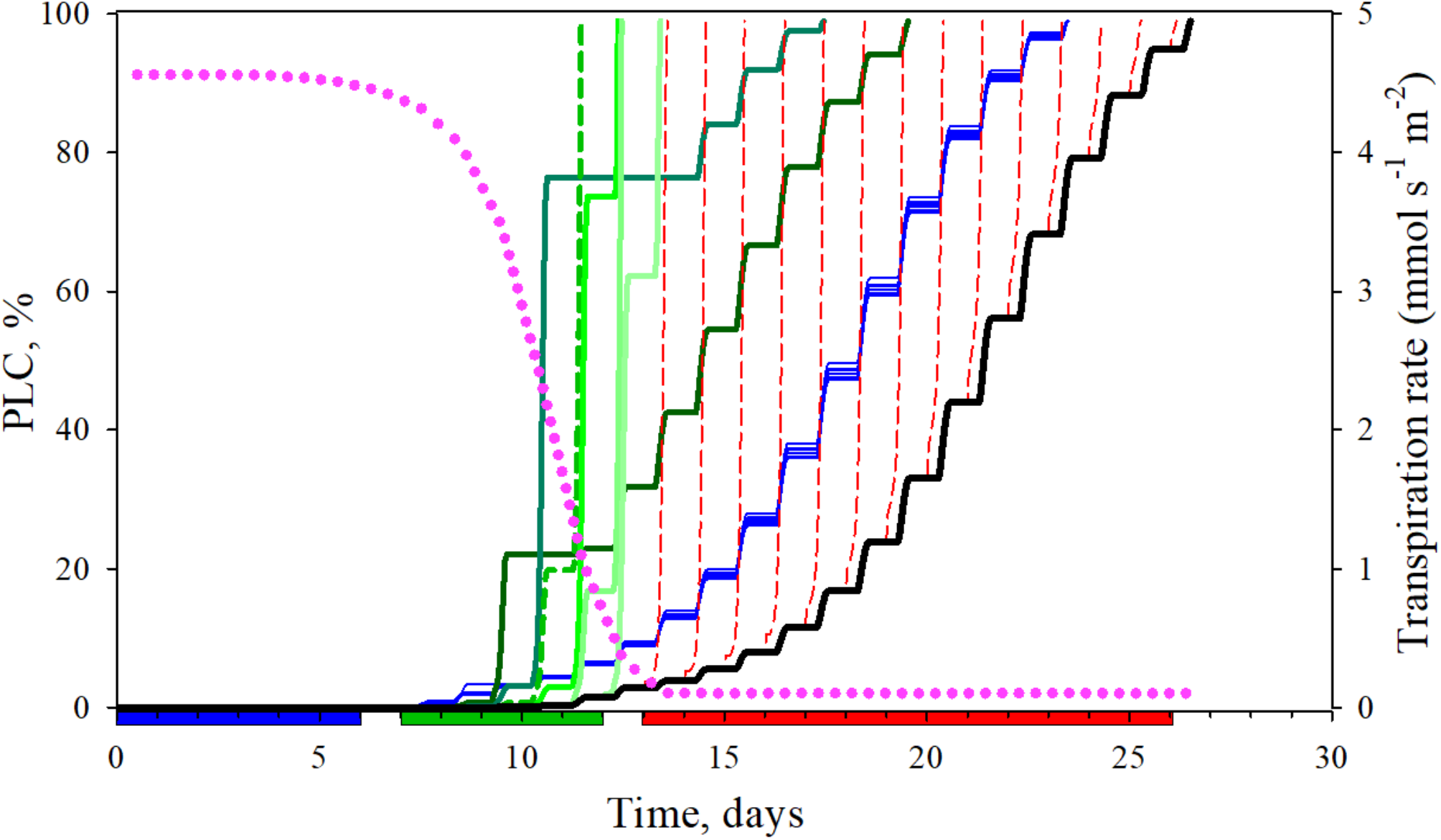
Simulated impacts of a heatwave on xylem embolism for a plant exposed to a water stress at day 0. The time courses of embolism (black line) and transpiration (dotted pink line) are shown for a control plant exposed only to water stress. The other lines show the time courses of embolism for plant exposed to a +15°C heatwave for 3 consecutive days occurring at different times during the drought event: between day 0 and day 6 (blue lines), between day 5 and day12 (green lines), after day 13 (red dashed lines).

When a heatwave occurred at the onset of the drought episode (between day 0 and day 6), *ie* when stomata were fully open, the impact was relatively minor and plant hydraulic failure occurred 3 days earlier (figure 5, red lines). This effect was explained by the transpiration increase during the heatwave and the earliest depletion on soil water content. By contrast, a heatwave occurring after the onset of stomatal downregulation to limit transpiration (day 7) had a great effect on the development of embolism and the timing of hydraulic failure (figure 5, green lines). At day 9 the impact on hydraulic failure was maximal with a *t*_*HF*_ reduced to 11 days (figure 3, dashed green line). Heatwaves arising when stomata are completely closed (after day 13, figure 5, dashed blue lines) provoked the total embolization of the xylem tissue within a few hours, strongly reducing the time to hydraulic failure compared to a control plant.

### The critical role of the leaf phase transition temperature

To explore this hypothesis further, I conducted a sensitivity analysis of the model in order to identify the most relevant traits associated with this risk of hydraulic failure. To this end, I simulated a 3 days, +15°C heatwave occurring at day 9, *ie* corresponding to the most critical situation from above. I then explored how *t*_*HF*_ responded to a variation of *Tp*, *Q*_*10a*_ and *Q*_*10b*_. The results are shown in figure 6 as 2D maps where *t*_*HF*_ values are displayed using a color spectrum. The symbol at the center of each graph correspond to reference simulation. The sensitivity analysis shows that increasing or lowering *Q*_*10a*_ and *Q*_*10b*_ by +-20% had only a marginal impact on *t*_*HF*_. By contrast, a small increase of the cuticle phase transition temperature *Tp* (*ca* 2°C) sharply increased *t*_*HF*_. This suggests that plant safety margin under these hot-drought conditions is determined by the difference between *Tp* and the maximum heatwave temperature *T*_*air-max*_.

**Figure 6:**
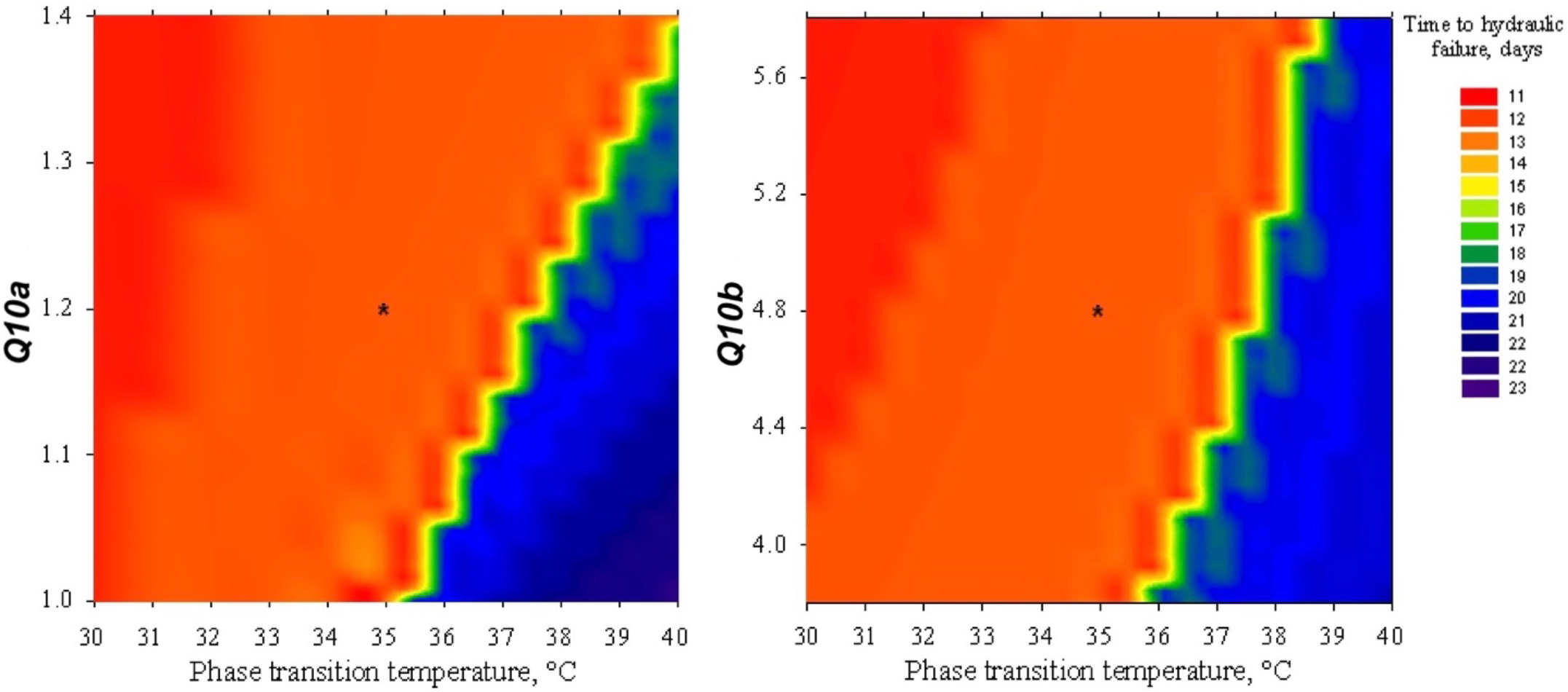
Effects of the phase transition temperature (*Tp*) and the *Q*_*10*_ of gmin below (*Q*_*10a*_) and above (*Q*_*10b*_) *Tp* on the time to plant hydraulic failure *t*_*HP*_. *t*_*HP*_ is shown as a color spectrum. The asterisks correspond to the control plant (*Tp = 35°C; Q_10a_ = 1.2; Q_10a_=4.8).*

To test this hypothesis, I analyzed the effects of *Tp* and *T*_*air-max*_ on *t*_*HF*_ (figure 7). The simulation shows that to maintain a constant safety margin (constant color in figure 7), *Tp* must be higher than *T*_*air-max*_ and increase linearly with *T*_*air-max*_ with a slope of *ca* 1.7 °C/°C. In other words, *Tp* must disproportionately increase when plant are exposed to hotter conditions. This is explained by the fact that leaf residual transpiration reaches critical values even when *T*_*air-max*_ is less than *Tp* because of the temperature effect on *g*_*min*_ and *VPD*.

**Figure 7:**
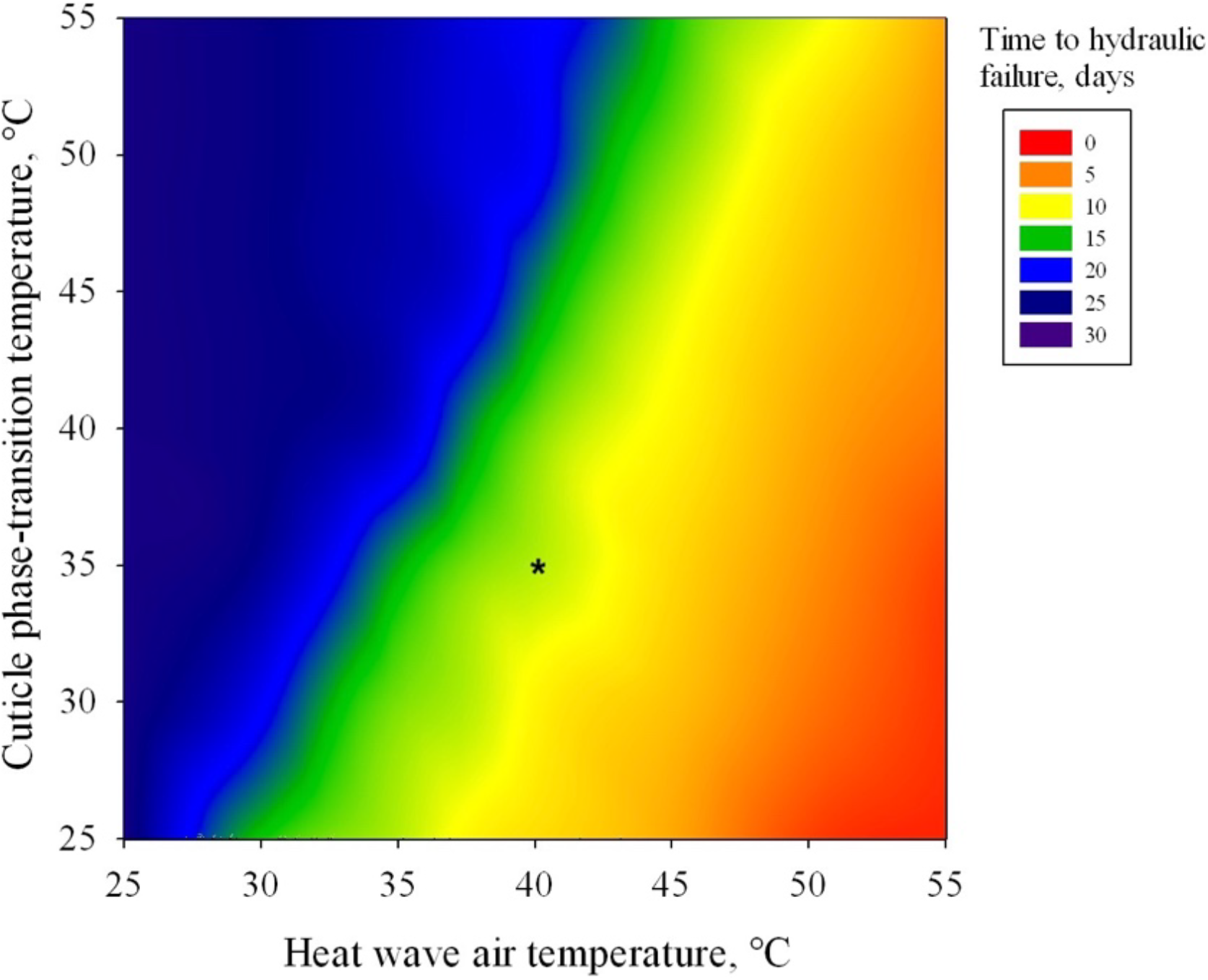
Effects of the phase transition temperature and the maximum heatwave temperature on the time to hydraulic failure. The asterisk corresponds to the control plant (*Tp = 35°C)* exposed to a +15°C heatwave.

## Discussion

As previously reported (Martin-StPaul *et al* 2017, Duursma *et al* 2019), the simulations of the mechanistic SurEau_C model suggest a central role of leaf residual transpiration on the kinetic of cavitation induction, the timing of plant hydraulic failure and mortality during a drought event. By integrating a new model of leaf cuticular conductance incorporating its bi-phasic temperature dependence, I propose here a new hypothesis to explain the striking increase in tree mortality under both hot and dry conditions. The causes of mortality in drought-prone trees are multiple and complex. Droughts may be the sole cause of tree mortality, but they are also often factors predisposing trees to lethal attacks by pathogen infections. During water stress, stomata close to prevent the deleterious formation of cavitation event in the xylem tissue (Tyree and Sperry 1989) but this comes at the cost of a reduced carbon uptake and a higher leaf temperature. Leaf water loss is then reduced to a residual value caused by water losses through the cuticle or leaky stomata. Because of the known exponential increase of the cuticular conductance *g*_*min*_ above a phase transition temperature *Tp* (Riederer and Muller 2006) and because of the high *VPD* associated with high air temperature, leaf residual transpiration increases sharply under hot conditions. Under these stringent conditions, plants cannot contain the formation of cavitation and a phenomenon of run-away embolism occurs leading very suddenly to a total hydraulic failure. This critical situation occurs only when plants are exposed to a water-stress intense enough to impact stomatal conductance and at the same time exposed to a heatwave. This putative process could provide a new mechanistic explanation for the mortality events associated with hot-drought events.

The model suggests a critical role of the cuticle phase transition temperature *Tp* on plant safety margin for hydraulic failure during hot-drought conditions. *Tp* is known to vary significantly across species (Riederer and Muller 2006) but the data available in the literature on this parameter is still very scarce and a correlation between *Tp* and species hot-drought resistance cannot be made. Quite strikingly, *Rhazya stricta*, a typical woody hot-desert plant, possesses a very high *Tp* value (>50°C) and also a greater proportion of triterpenoids in its cuticle (Schuster *et al* 2016). This may suggest that plant could achieve a higher cuticular thermostability by modifying the chemical composition of their cuticles but the degree of genetic variability or plasticity of *Tp* is not yet known.

This paper points to the importance of considering *Tp* as a key trait when evaluating hot-drought tolerance in plants. Other key functional traits, like xylem cavitation resistance *P*_*50*_ or plant hydraulic capacitance *C,* are not known to display such a temperature dependence, emphasizing the putative role of *g*_*min*_ in plant hot-drought resistance. There is very little work on the plasticity of *g*_*min*_ and especially of *Tp* (Duurema *et al* 2019). If the importance of these traits for the survival of species in heat drought is confirmed, it would be important to explore the acclimatization capacities of trees for these traits.

My conclusions are based on model simulations and experimental confirmations are urgently awaited. For this, it is first necessary to accurately estimate the value of *Tp* for different species. We have recently proposed a new experimental set-up to measure this parameter (Billon *et al* 2020). Then, the objective is to measure the speed of embolism formation for plants exposed to temperatures below and above *Tp* and variable degrees of water stress. The optical method recently developed by Brodribb *et al* (2016) is very appropriate for such measurements. Cavitation is expected to form more rapidly for plants that are stressed and exposed to temperatures above *Tp*. The experiment can easily be done on cut twigs or on potted plants. Validating the predictions of the model *in situ* is more challenging. Tree transpiration flux detection methods are probably not sensitive enough to detect the increase in cuticle water loss beyond *Tp* on stressed trees, but it would be interesting to explore these experimental data with this new look. A very different approach would be to analyze the emissions of volatile organic compounds at the ecosystem scale (Niinemets and Monson, 2013). The increased permeability of the cuticle to water at high temperatures could also result in a higher emission of these compounds.

I hope that this paper will stimulate new researches to explore further the role of *g*_*min*_ and *Tp* on plant hot-drought resistance. New tools for the high throughput phenotyping of these parameters will have to be developed in order to identify the putative underlying genes and help breeders to screen for genotypes better adapted to future climatic conditions.

## Supplementary material

The C code of the SurEau program is available from the data INRAE public depository: https://data.inrae.fr/dataset.xhtml?persistentId=doi:10.15454/6Z1MXK

## Acknowledgements

I was supported in part by IDEX-ISITE initiative 16-IDEX-0001 (CAP2025) and the ANR-18-CE20-0005 *Hydrauleaks*. Version 2 of this preprint has been peer-reviewed and recommended by Peer Community In Forest and Wood Sciences (https://doi.org/10.24072/pci.forestwoodsci.100002).

## Appendix

## Appendix 1: List of abbreviations

*e*_*leaf*_ (kPa): Vapor pressure of water in the leaf at *T*_*leaf*_ and ⍰_*leaf*_
*e*_*sat_leaf*_ (kPa): Saturation vapor pressure in the leaf at *T*_*air*_
*e*_*air*_ (kPa): Vapor pressure of water in the air at *T*_*air*_
*e*_*sat_air*_ (kPa): Saturation vapor pressure of the air at *T*_*air*_
*E* (mmol s^−1^ m^−2^): Leaf transpiration rate
*F*_*T*_ (poise^−1^): Dynamic fluidity of water
*g*_*leaf*_ (mmol s^−1^ m^−2^): Maximum leaf conductance to vapor
*g*_*min*_ (mmol s^−1^ m^−2^): Minimum (residual) leaf conductance
*g*_*s*_ (mmol s^−1^ m^−2^): Leaf stomatal conductance
*K*_*plant*_ (mmol s^−1^ MPa^−1^): Whole plant hydraulic conductance
*K*_*T*_ (mmol s^−1^ MPa^−1^): Hydraulic conductance
*P*_*air*_ (kPa): Atmospheric pressure
*P*_*leaf*_ (MPa): Leaf turgor pressure
*P*_*50*_ (MPa): Xylem pressure inducing 50% loss of hydraulic conductance
*Q*_*10a*_: Leaf cuticle *Q10* value below *Tp*
*Q*_*10b*_: Leaf cuticle *Q10* value above *Tp*
*RH*_*air*_ (%): Air relative humidity
*RH*_*air-min*_ (%): Minimum daily air relative humidity
*RH*_*air-max*_ (%): Maximum daily air relative humidity
*t*_*g0*_ (s): Time to stomatal closure
*t*_*HF*_ (s): Time to hydraulic failure
*T*_*air*_ (°C): Air temperature
*T*_*air-min*_ (°C): Minimum daily air temperature
*T*_*air-max*_ (°C): Maximum daily air temperature
*T*_*leaf*_ (°C): Leaf temperature
*Tp* (°C): Leaf cuticle phase transition temperature
*VPD* (kPa): Vapor pressure deficit between the leaf and the air
*ε* (MPa): Leaf modulus of elasticity
*σ*_*T*_ (Nm_−1_): Water surface tension.
*Ψ*_*leaf*_ (MPa): Leaf water potential
*Π*_*T*_ (MPa): Leaf osmotic potential

